# A histidine kinase gene is required for large radius root tip circumnutation and surface exploration in rice

**DOI:** 10.1101/437012

**Authors:** Kevin R. Lehner, Isaiah Taylor, Erin N. McCaskey, Rashmi Jain, Pamela C. Ronald, Daniel I. Goldman, Philip N. Benfey

**Affiliations:** Department of Biology, Duke University, Durham, NC 27708; University Program in Genetics and Genomics, Duke University, Durham, NC 27708; School of Physics, Georgia Institute of Technology, Atlanta, GA 30332; Department of Plant Pathology and the Genome Center, University of California, Davis, CA 95616; Howard Hughes Medical Institute, Duke University, Durham, NC 27708

## Abstract

The intricate growth patterns that accompany plant organ elongation have long intrigued biologists ^1^. Circumnutation refers to the circular or elliptical growth of the tip of a plant organ around a central axis. While the utility of circumnutation for climbing plants is clear, its function in roots is less obvious. Additionally, the genetic requirements for root circumnutation are not known. Here we show that mutations in a gene encoding a histidine kinase abolish large radius root tip circumnutation in rice. Using a gel-based imaging system and a whole genome sequenced mutant population, we identified three different mutant alleles of the gene *OsHK1* that exhibit increased seedling root depth. Time-lapse imaging indicated that this phenotype is likely due to a lack of large radius root tip circumnutation in *OsHK1* mutants. Treatment of mutant roots with the plant hormone zeatin rescues circumnutation, indicating that *OsHK1* functions in a cytokinin-related signaling pathway. We found that *OsHK1* mutants are impaired in their ability to explore flat surfaces, suggesting that circumnutation facilitates root exploration at the interface of compacted soil horizons.

---

In a classic work, Charles and Francis Darwin measured the growth movements of shoots and roots across a wide variety of species ^1^. When grown between two smoke-covered glass plates, the roots of some species showed traces suggesting root tip circumnutation. This root growth pattern has since been described in multiple plant species ^2–4^. The extent to which these movements represent sustained oscillatory processes or more random patterns remains under debate ^5^. Root circumnutation has been suggested to reduce soil penetration costs in maize ^6^ and to facilitate seedling establishment in rice ^7^. In *Arabidopsis thaliana*, when seedlings are grown on tilted agar plates, roots create a waving and skewing growth pattern. The mechanisms underlying these phenotypes have been attributed to a complex interplay of factors, including gravitropic, thigmotropic, and physical surface responses ^8^. It has been proposed that circumnutation functions in generating and modulating these patterns ^9^.

A small number of genes have been identified that are required for shoot circumnutation ^10–12^. However, the genetic control of these processes appears to differ between shoots and roots. For example, while mutations in the *LAZY1* gene abolish coleoptile circular growth, the gene is not required for root tip circumnutation in rice ^13^.

Root system architecture (RSA) describes the spatial arrangement of roots within a growth substrate, which is a function of the lengths, branching patterns, angles, and numbers of roots, among other factors ^14^. Here we identify mutations in a histidine kinase gene that profoundly affect RSA through the alteration of root circumnutation. While measuring RSA in cereals has historically focused on static traits, the growth dynamics of roots is likely a critical parameter contributing to root architecture. Two recently cloned genes that affect root growth angle in cereals have roles in regulating root gravitropism ^15,16^. The study of the interplay between root growth behavior and RSA is a rapidly developing field.

Using a gel-based imaging system ^17^ we measured RSA phenotypes of lines derived from a sequence-indexed fast-neutron irradiated mutant rice population in the Kitaake background ^18^. Kitaake is a rapid cycling, largely photoperiod insensitive Japonica cultivar. From this analysis, we identified segregants within a mutant line, FN-287, which had increased early root depth when compared to its wild-type parent (Fig. 1a). This phenotypic difference appeared to be driven by a much deeper primary root. We identified a single base substitution that cosegregated with the deep root phenotype. The SNP is predicted to result in a premature stop codon in exon 1 of *OsHK1*, a gene encoding a putative histidine kinase (HK) ^19^. We identified two additional lines that harbor likely loss-of-function mutations in *OsHK1* (Extended Data Fig. 1). All three confirmed homozygous mutants had root depths that were twice that of their wild-type parent (Fig. 1b). An F_1_ derived from a cross of two of the mutants (*oshk1-1* x *oshk1-2*) failed to show complementation of the mutant phenotype (Fig. 1a). We conclude that the deeper root RSA phenotype is attributable to mutations in *OsHK1*.

**Figure 1:**
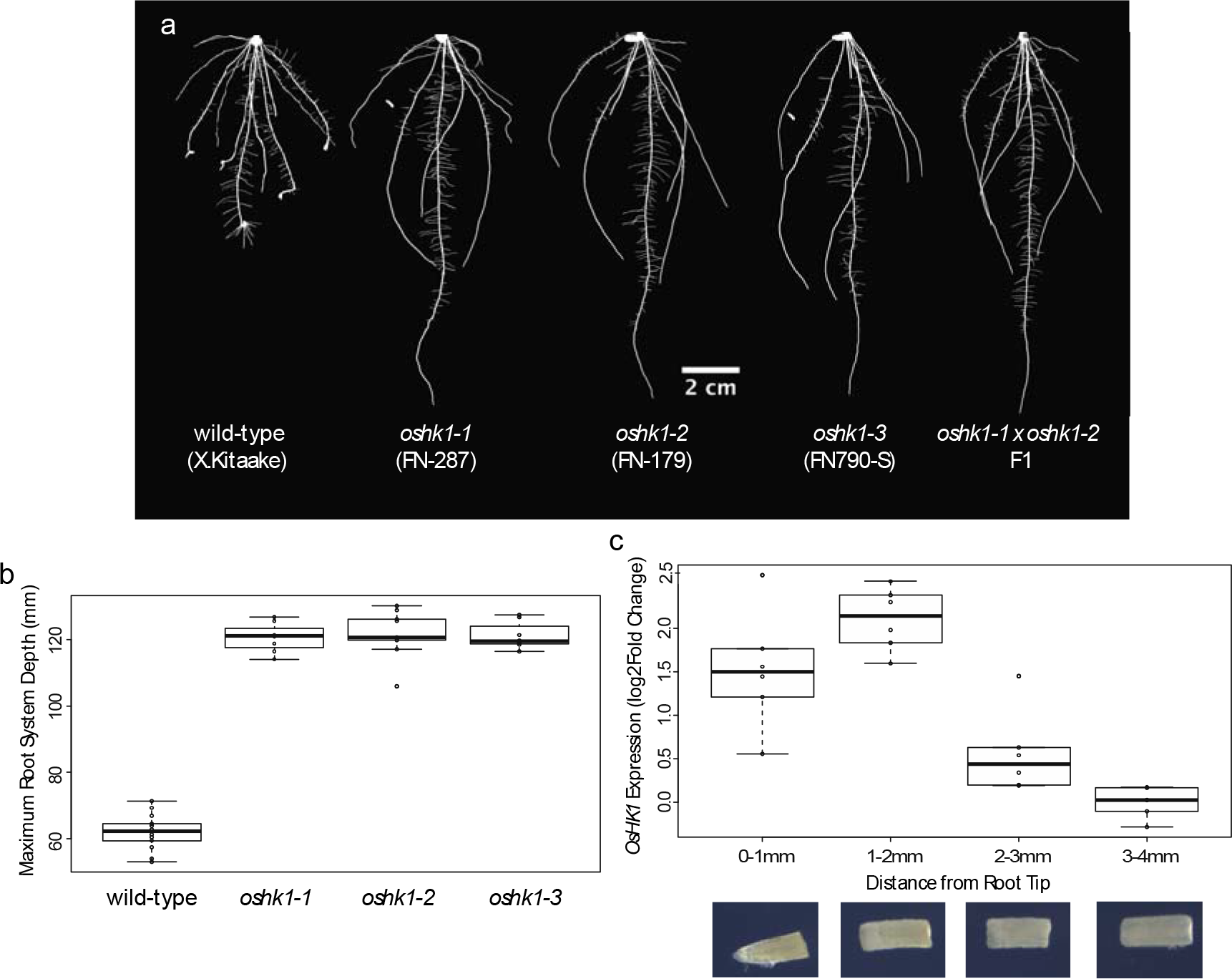
*OsHK1* mutants have increased root depth. **a,** Thresholded images of seven-day old seedlings grown in Yoshida’s nutrient media, solidified with 0.25% gellan gum. **b,** Quantification of Network depth from GiA Roots. Value for each plant is the mean of 40 rotational images. In the boxplots, the thick middle line represents the median, the upper and lower edges of the box denote the first and third quartile, respectively. Wild-type differs from all three *OsHK1* mutants at p<2E-16. **c,** qPCR expression level of *OsHK1* in wild-type roots. *OsHK1* expression differs between the two sections proximal to the tip (0-1, 1-2mm) as compared to the two sections distal to the tip (2-3, 3-4mm) at p<0.01.

HKs were identified as members of two-component signaling pathways that facilitate environmental sensing in bacteria ^20^. In modified forms, these molecules also play important roles in plants ^21^, particularly as receptors for the hormone cytokinin ^22^. Along with a cytokinin-binding CHASE domain, these proteins have a conserved histidine residue in the kinase domain and an aspartate residue in the receiver domain. *OsHK1* encodes a HK that is structurally related to cytokinin receptors, yet is lacking the CHASE and transmembrane domains (Extended Data Fig. 2). *OsHK1* has high similarity to Arabidopsis *AtCKI2/AtAHK5*, which has been implicated in diverse processes such as stomatal signaling, pathogen response, and hormone-induced root elongation ^23–25^.

In mature plant organs, *OsHK1* is most highly expressed in roots (Extended Data Fig. 3). To determine where along the root *OsHK1* is expressed, we performed qRT-PCR on dissected 1mm sections of wild-type seedlings and found the highest expression in the section 1-2 mm from the root tip (Fig. 1c). This region is within the elongation zone of the root ^26^.

To better understand the increased root depth phenotype in *OsHK1* mutants, we imaged primary root growth in gel-based media at 15-minute intervals for 2-3 days after emergence. We observed a striking difference in root tip circumnutation between wild-type (Fig. 2a, Supplementary Movie 1, Extended Data Fig. 4) and *OsHK1* mutants (Fig. 2b, Supplementary Movie 2, Extended Data Fig. 4). During very early stages of primary root elongation, growth was similar between wild-type and mutant plants. Subsequently, wild-type primary roots began entrainment of an oscillatory pattern leading to large radius root tip circumnutation (Fig. 2c), which did not occur in the mutants (Fig. 2d). We calculated the amplitude of circumnutation as the lateral displacement of the tip relative to the center of the root, analogous to the radius of a cylinder circumscribed by the root tip. Wild-type root tip circumnutation reached a maximum amplitude of 0.8 mm (Extended Data Fig. 5a). The period of circumnutation increased over time in wild-type roots, with a maximum period of 3.5 hours (Extended Data Fig. 5c). The lack of large diameter root tip circumnutation is likely the major factor contributing to the deep root phenotype that we identified in the *OsHK1* mutants.

**Figure 2:**
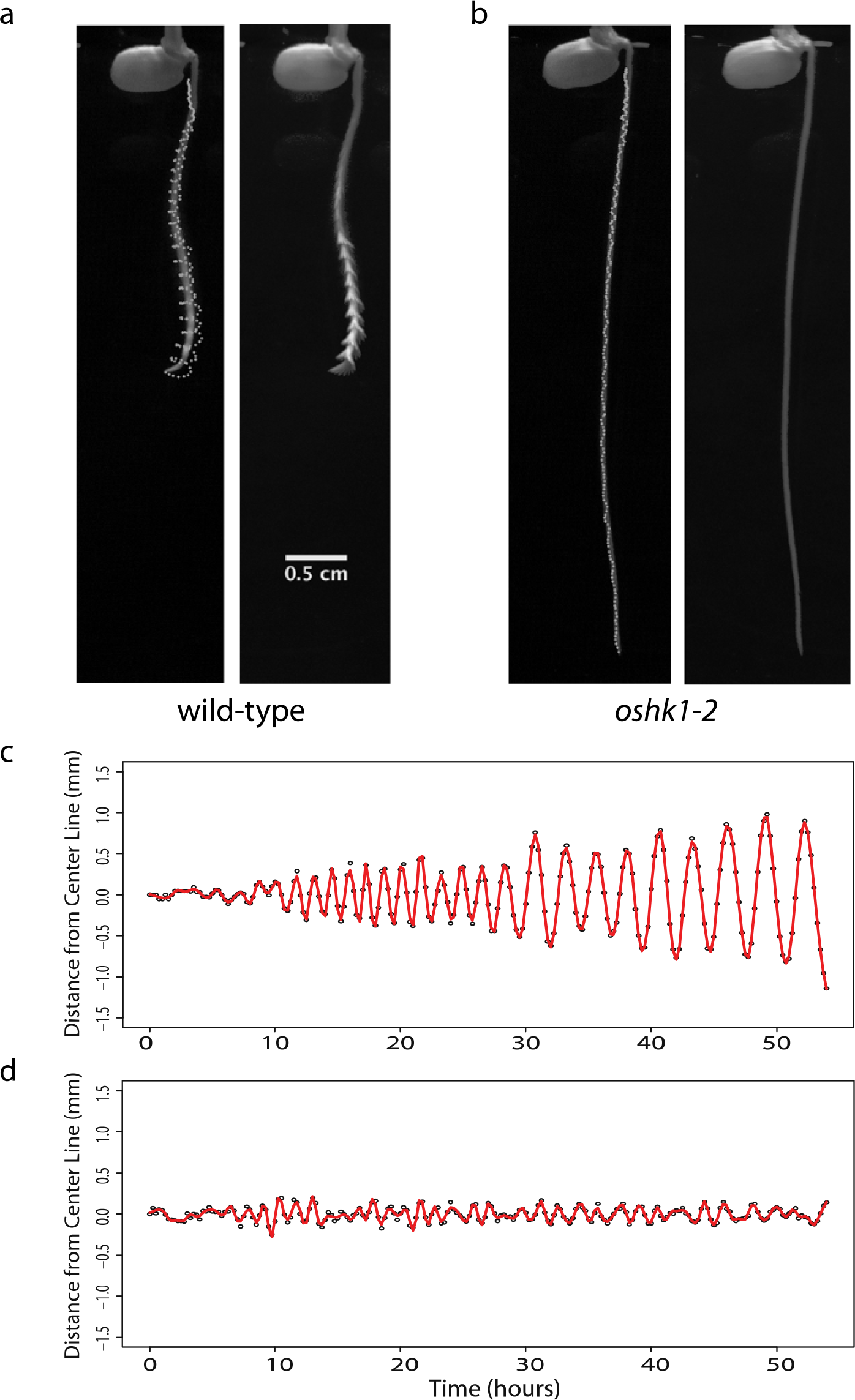
*OsHK1* is required for large radius root tip circumnutation. Tip position at 15-minute intervals over 54 hours of imaging is shown as points (left), and as maximum intensity projections (right) for **a,** wild-type, and **b,** *oshk1-2*. Plots of tip position relative to the midline of the root for **c,** wild-type, and **d,** *oshk1-2*. The red line represents a fitted natural cubic spline.

Circumnutatory processes, such as that in some vining plants, are often characterized by a fixed direction of rotation, or handedness ^27^. We found no evidence for a strong pattern of handedness in the roots we observed. From a sample of 21 wild-type roots, 11 showed a clockwise rotation and 10 circumnutated in a counterclockwise direction (Extended Data Fig. 8).

After 48 hours of growth, the tips of some wild-type roots began to wind over themselves, forming a knot-like structure (Supplementary Movie 3). This is similar to the “Root Meander Curling (RMC)” phenotype previously described in rice ^28,29^. In our gel imaging experiments, we found evidence of RMC-like behavior in all wild-type replicates, but this phenotype did not occur in any of the *OsHK1* mutants (Extended Data Fig. 9). This suggests an integral link between root tip circumnutation and the RMC trait. In a first attempt to determine the extent of conservation of circumnutation, we examined four diverse rice cultivars and found patterns of root tip circular movement in all of them (Extended Data Fig. 10), suggesting that this phenotype may be a general property of the species.

To investigate the molecular signaling mechanisms regulating circumnutation, we performed RNA-Seq on wild-type and mutant tissue spanning 1 to 2 mm from the root tip, where *OsHK1* is maximally expressed. Differential expression analysis identified 622 genes higher and 119 genes lower in wild-type as compared to mutant (Extended Data Table 1). Because HKs are known to regulate hormone signaling pathways, we performed GO term analysis and screened for terms associated with hormone signaling. We found enrichment for *two-component signal transduction system (phosphorelay)* and *two-component response regulator activity*, closely associated with canonical cytokinin signal transduction (Extended Data Table 2). We also identified 8 cytokinin responsive Type-A Response Regulator genes ^30^ with detectable expression in this region of the root. Six of these displayed statistically significant reduction in expression in the mutant (Fig. 3a). We additionally observed highly significant overlap between genes previously demonstrated to be regulated by cytokinin treatment in rice roots and the differentially expressed genes identified from our RNA-Seq data ^31^ (Fig. 3b). Taken together, these results indicate *OsHK1* positively regulates a cytokinin-related signal transduction pathway.

**Figure 3:**
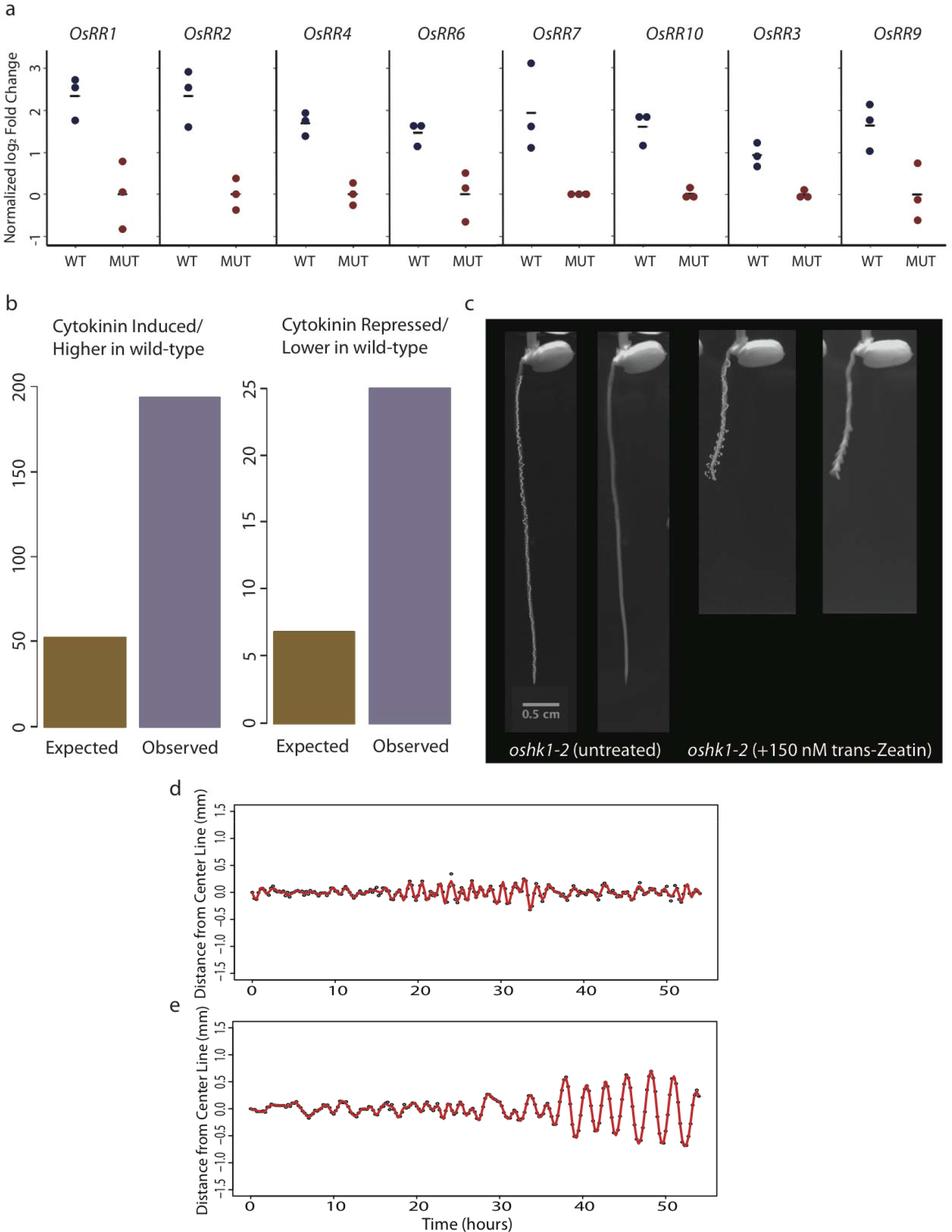
*OsHK1* controls circumnutation through activation of a cytokinin-related signaling pathway. **a,** RNA-seq data of normalized levels of expressed Type-A Response Regulator genes from root sections 1-2mm proximal to the tip. Mutant points represent individual replicates from all three *oshk1* mutant alleles. Expression levels of *OsRR1, 2, 4, 6, 7,* and *10* are significantly reduced at a FDR <0.05. **b,** Enrichment analysis of differentially expressed genes from this study with cytokinin responsive genes previously identified in rice roots. Left chart compares expected and observed overlap of cytokinin-induced genes and those higher in wild-type than mutant. Right chart compares expected and observed overlap between cytokinin-repressed genes and those lower in wild-type than mutant. p<.0001 for both comparisons, hypergeometric test. **c,** Summary of 54 hours of imaging (as in Fig. 2) for *oshk1-2,* untreated (left two panels) and treated with 150 nM trans-zeatin (right two panels). Plots of tip position (as in Fig. 2) for *oshk1-2* (**d**), and *oshk1-2* with 150 nM trans-zeatin (**e**).

Based on these data, we hypothesized that *OsHK1* acts through a downstream pathway related to cytokinin response, but *OsHK1* itself has lost its ability to respond to cytokinin. This raised the possibility that loss of *OsHK1* activity could be compensated by activation of the canonical cytokinin signaling pathway. To test this hypothesis, we germinated mutant seeds in media containing a concentration range of the naturally occurring cytokinin, trans-zeatin. For concentrations at or above 80 nM we observed robust rescue of circumnutation in the mutant (Fig. 3c, Extended Data Fig. 11). This result is consistent with our hypothesis that loss of *OsHK1* can be rescued by the activation of latent canonical cytokinin signaling processes. Overall these data indicate that rice root circumnutation is controlled by *OsHK1*-mediated activation of a cytokinin-related signaling pathway.

In light of the conserved nature of rice root tip circumnutation, we sought to understand possible functions of this pattern of growth. In highly compacted soil horizons, wheat roots have been shown to primarily utilize cracks and biopores to penetrate into deeper strata ^32^. Upland and rainfed lowland rice are similarly affected by hardened subsoil layers ^33^. While a number of root traits have been suggested to improve tip penetration in conditions in which soil mechanical impedance is a limiting factor in growth ^34^, little is known about the roles of circular root tip growth at the interface of compacted soil horizons.

We tested the hypothesis that circumnutation is required to facilitate effective surface exploration by roots. This was motivated by a phenotype that we observed in *OsHK1* mutants, in which coils were formed when roots encountered flat surfaces (Fig. 4a). We hypothesized that this coiling behavior of *OsHK1* mutants would inhibit surface exploration when roots encountered a flat surface such as a compacted soil horizon.

**Figure 4:**
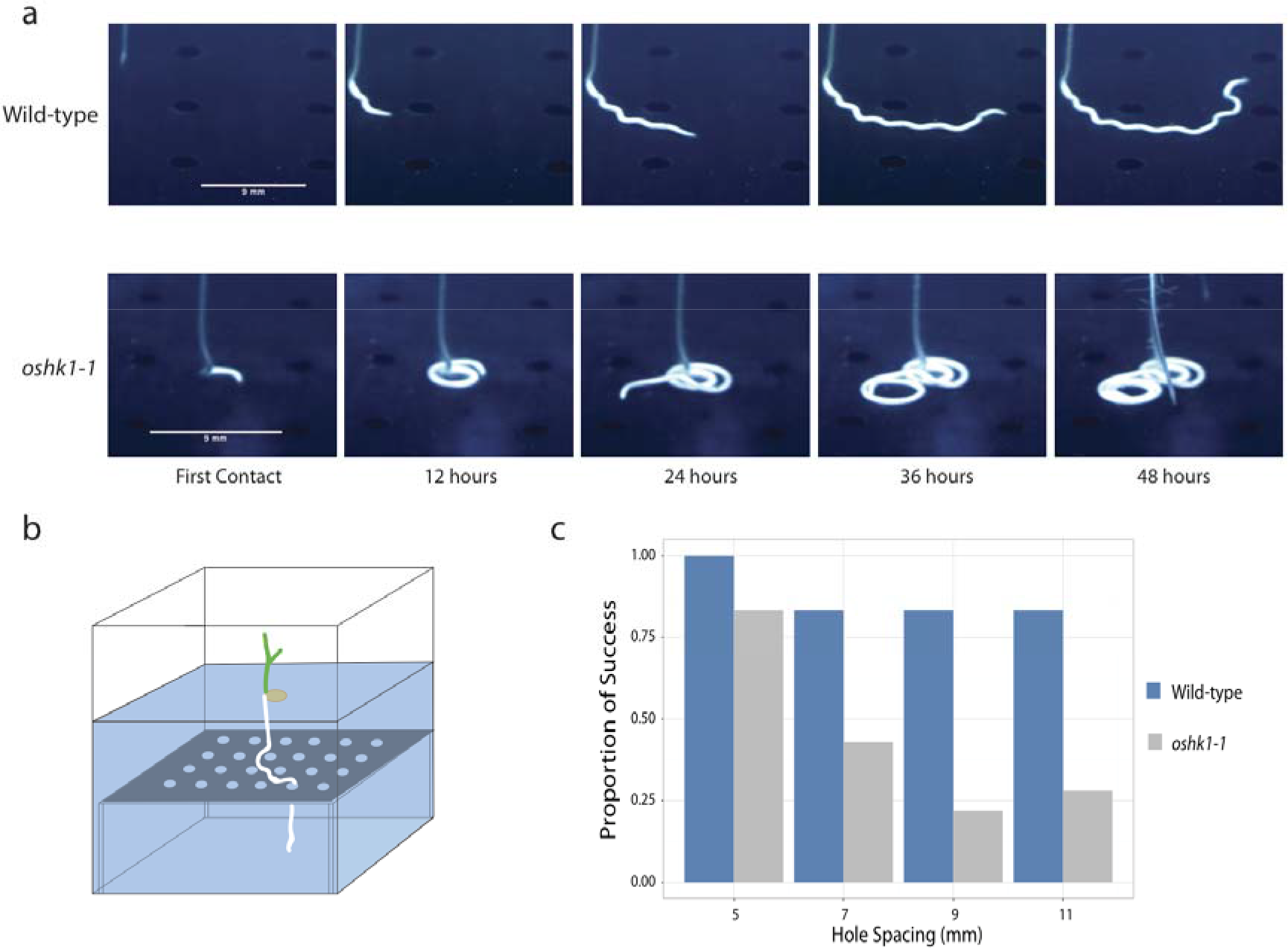
*OsHK1* is required for effective root tip surface exploration. **a,** Time lapse images showing phenotype of wild-type and mutant after encountering flat surface. **b,** Example of root surface exploration assay. **c,** Proportion of success in finding holes arranged in a square pattern on a flat platform after 72 hours across 4 different hole spacings. n = 11, 12, 6, and 6 trials for wildtype and 12, 16, 18, and 7 trials for mutant at 5, 7, 9, and 11 mm hole spacings, respectively.

We modeled this environment systematically using plastic surfaces with holes of 2.5mm diameter equally spaced in a square lattice at different densities. The surfaces were placed on hollow platforms of equal height in containers with gel-based media (Fig. 4b). A high-throughput automated imaging system acquired images of growing roots from two cameras at 15-minute intervals. One camera was positioned facing the front of the growth container to visualize root tip circumnutation and root depth, while the second was placed at an angle to capture the behavior of the root along the surface.

At the highest hole density (5mm spacing), both wild-type and *OsHK1* mutant primary roots were effective in finding holes and continuing deeper growth (Fig. 4c). However, as hole density decreased, *OsHK1* mutant roots showed a reduction in success in finding a hole. To quantify this effect, we employed logistic regression to model the probability of success in finding a hole as a function of spacing and genotype (Extended Data Fig. 12). We observed a statistically significant effect of genotype translating to an overall estimated 10.6-fold increase in the odds of success in the wild-type compared to mutant (estimated odds ratio 10.59, 95% CI (3.35, 42.63)).

These data indicate that wild-type roots are more effective in exploration and less affected by sparse hole density than *OsHK1* mutants, providing a plausible mechanism to buffer against environmental uncertainty inherent in soil surface exploration. We propose circular tip movement provides a mechanism to break the intrinsic root coiling pattern seen in *OsHK1* mutants, which would otherwise restrict exploration to spatially confined surface domains, and that this movement consequently promotes root exploration (Fig. 4a). Successful navigation through cracks or biopores is likely to allow for more effective penetration into deeper soil strata.

In summary, our work emphasizes the intimate link between the dynamics of root growth and root architecture, and suggests the possibility of modulating circumnutation to better allow roots to penetrate compacted soil horizons.

## Acknowledgments

The authors wish to thank the Duke Phytotron staff and Niba Nirmal for technical support; Mawsheng Chern and Guotian Li for helpful discussions provided during this project; Joe Kieber for helpful discussions; Erin Sparks, Colleen Drapek, and members of the Benfey Lab for critical reading.

## Author contributions

KRL and PNB conceived of this study. KRL, IT, ENM, RJ, PCR, DIG, and PNB designed methodology, performed experimental investigations, and analyzed data. KRL, IT, ENM, and PNB wrote the original drafts of this paper, with review provided by PCR and DIG.

## Extended Figures

**Extended Figure 1:**
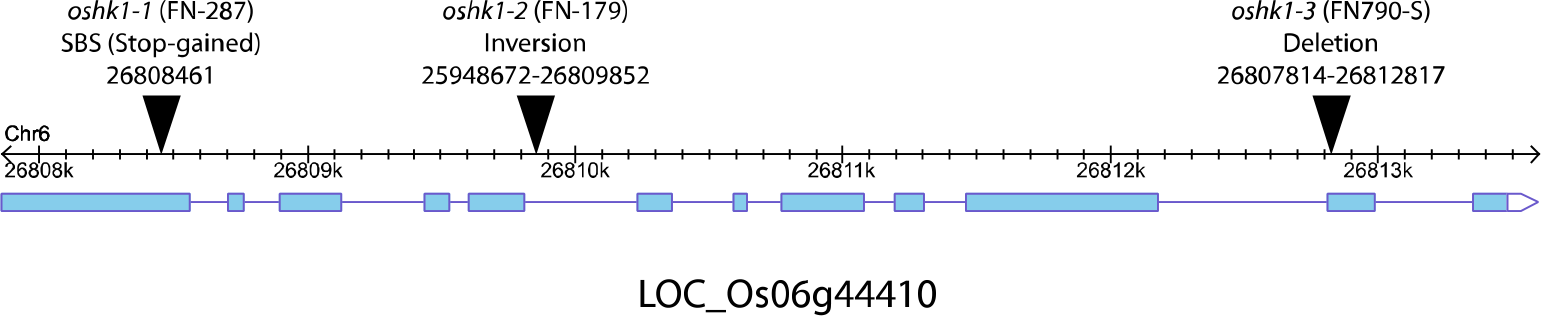
Gene model of *OsHK1* (LOC_Os06g44410) ^1^ including locations of exons (light blue). Positions of endpoints within gene in *oshk1* mutants are shown as black triangles.

**Extended Figure 2:**
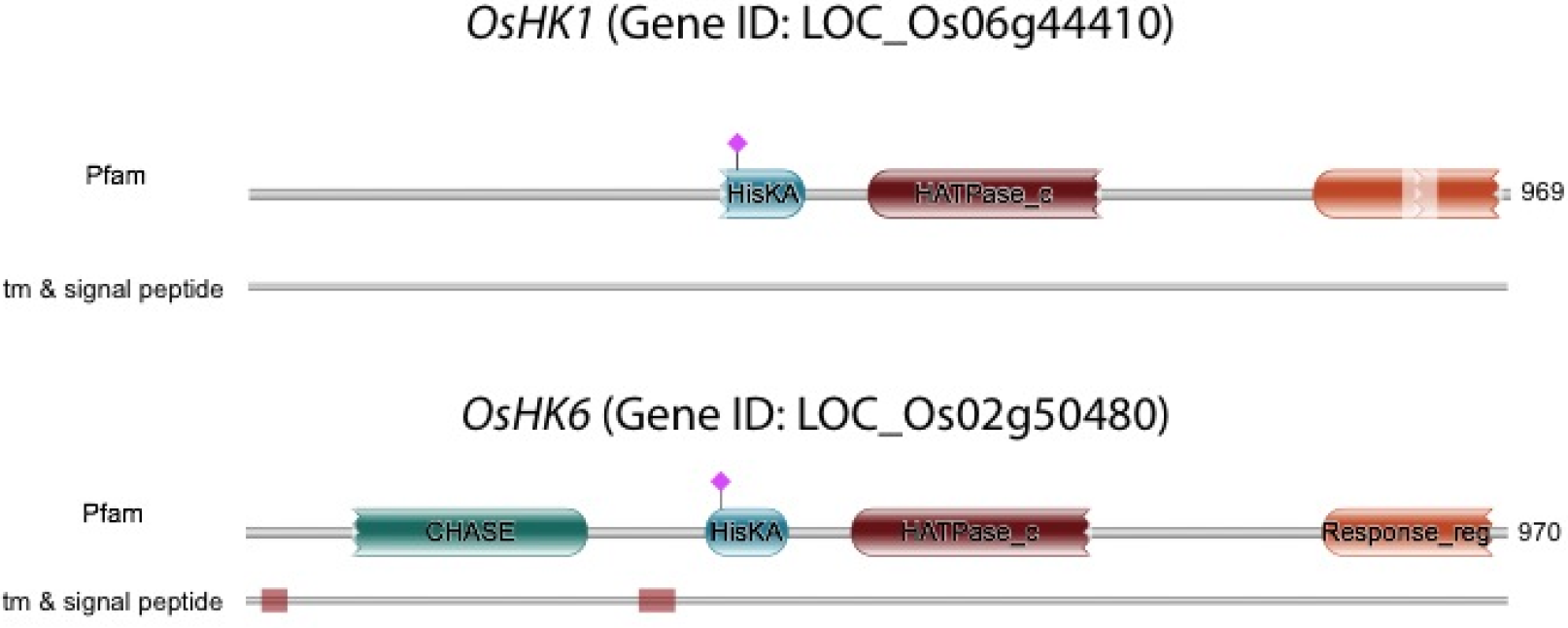
Protein domains of *OsHK1* and *OsHK6* predicted from HMMER ^2^. Location of predicted conserved histidine phosphorylation residues are shown as pink diamonds.

**Extended Figure 3:**
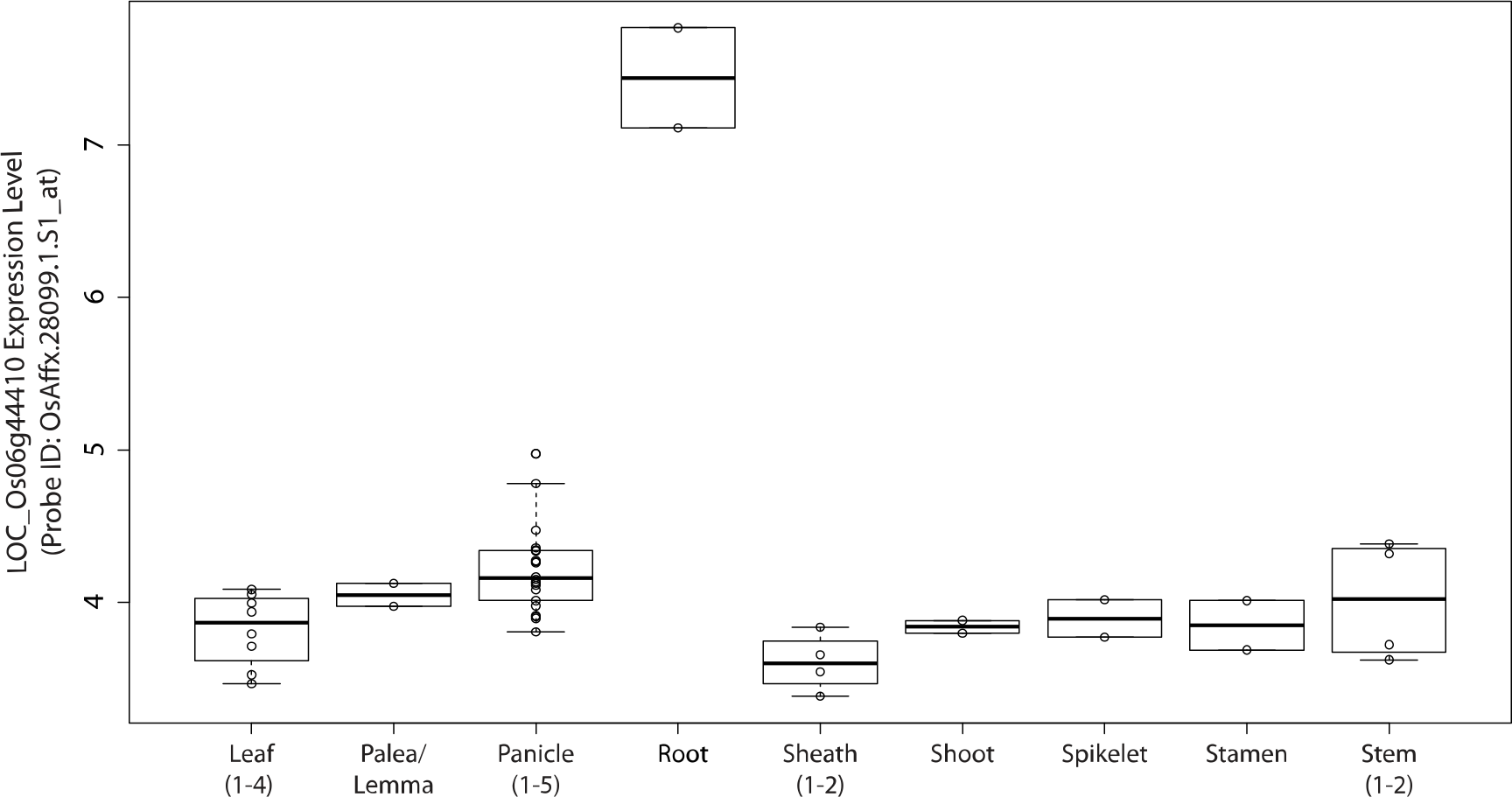
*OsHK1* expression level across mature organs in cultivar Minghui 63 ^3^. Plotted are normalized microarray expression values of probe ID: OsAffx.28099.1.S1_at from NCBI SRA BioProject PRJNA120617.

**Extended Figure 4:**
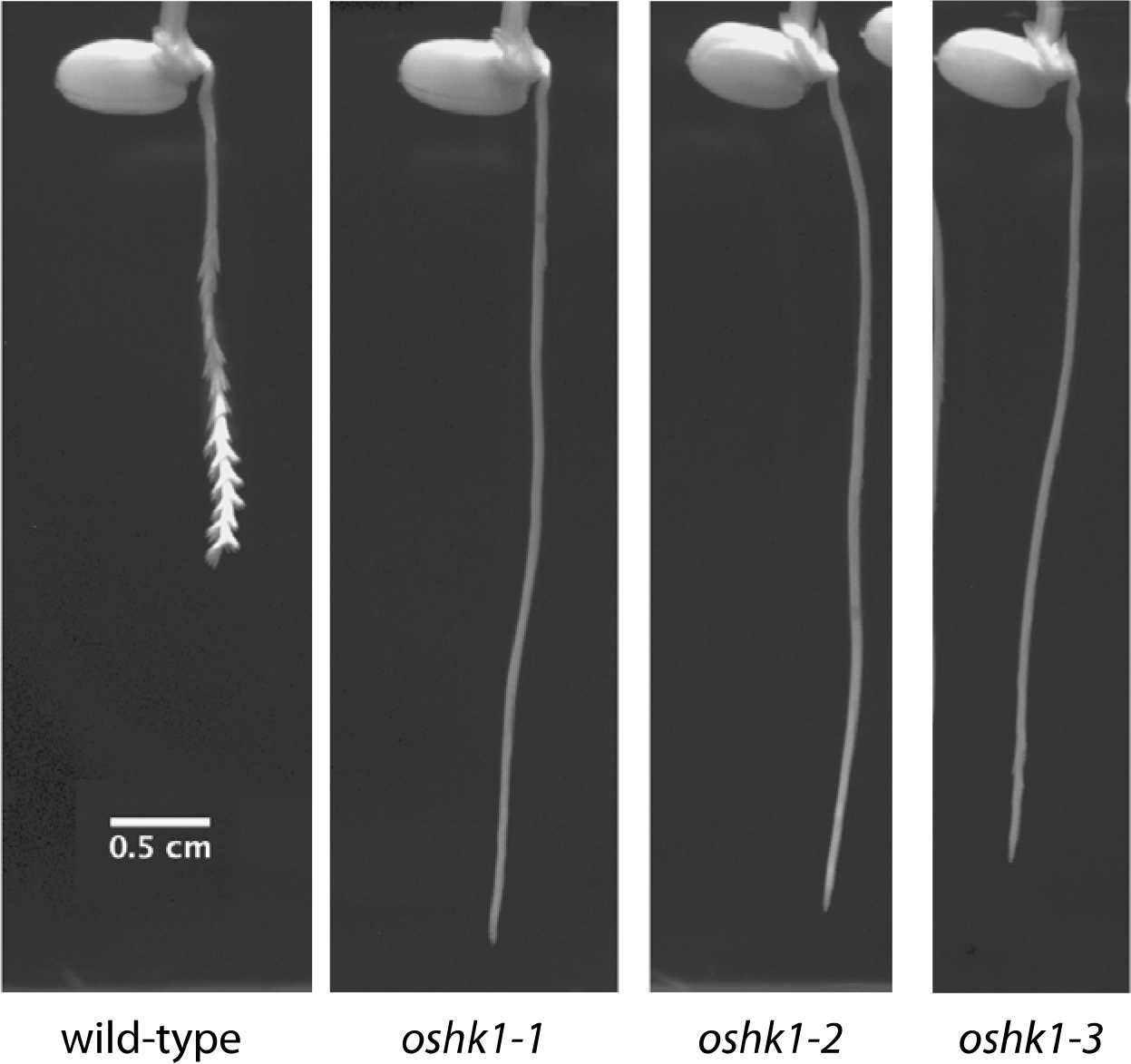
Maximum intensity projections of wild-type and *oshk1* alleles over 54 hours of growth, imaged at 15-minute intervals.

**Extended Figure 5:**
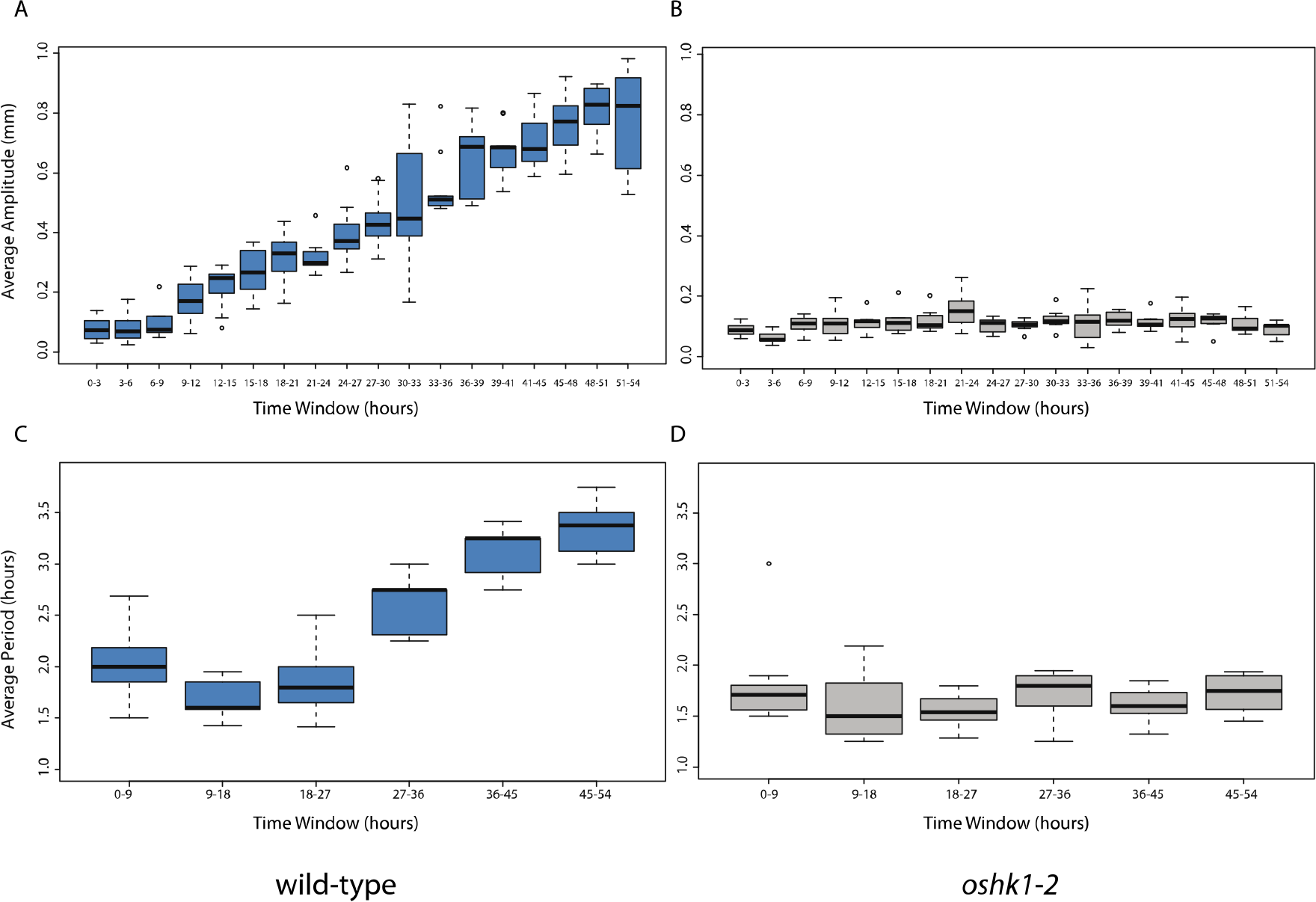
Root tip amplitudes (in millimeters) over 54 hours of growth for a, wild-type (n=9) and b, *oshk1-1* (n=7). Root tip periods (in hours) over 54 hours of growth for c, wild-type (n=9) and d, *oshk1-1* (n=7).

**Extended Figure 6:**
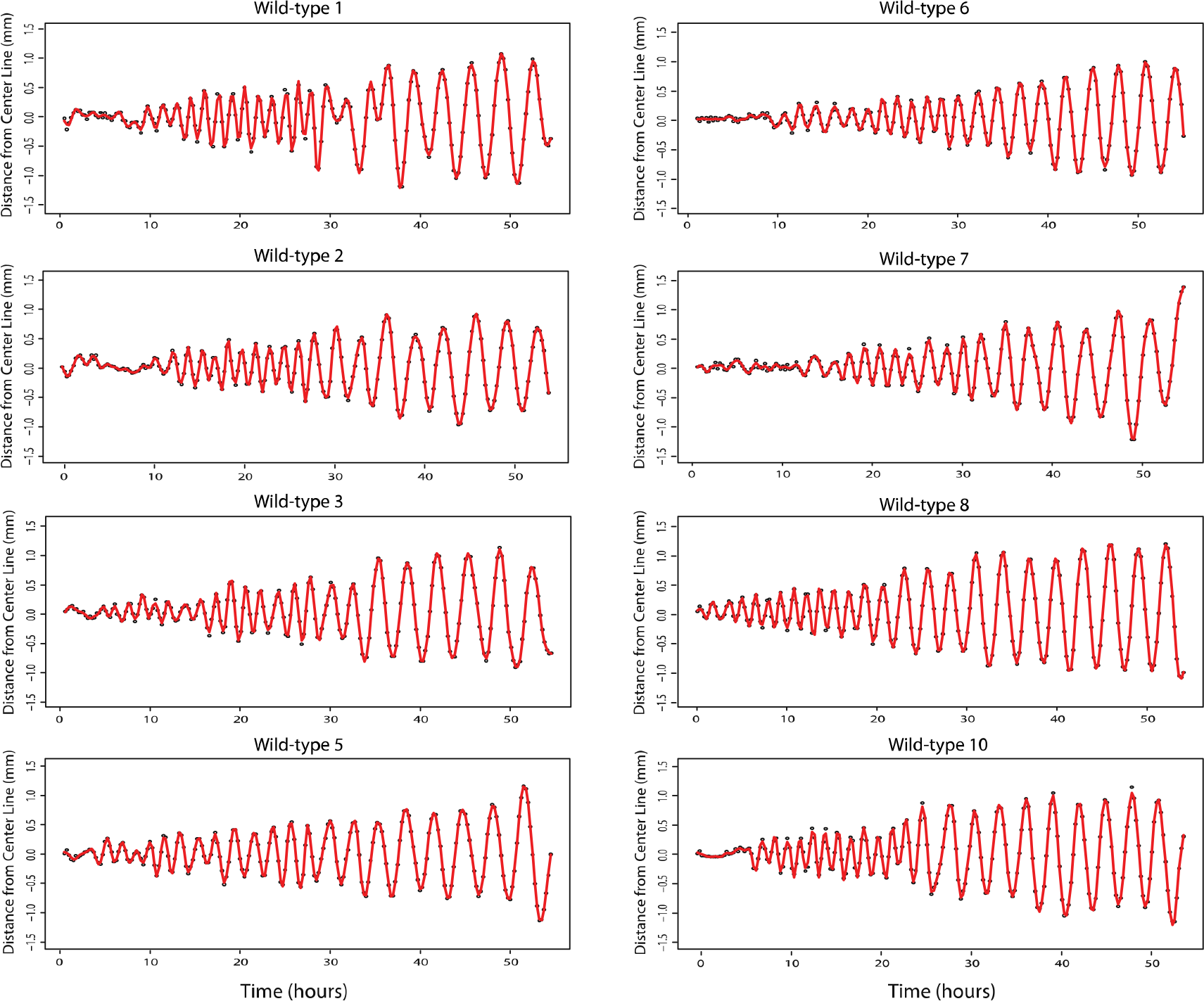
Additional plots of tip position relative to the midline of wild-type roots throughout 54 hours of growth.

**Extended Figure 7:**
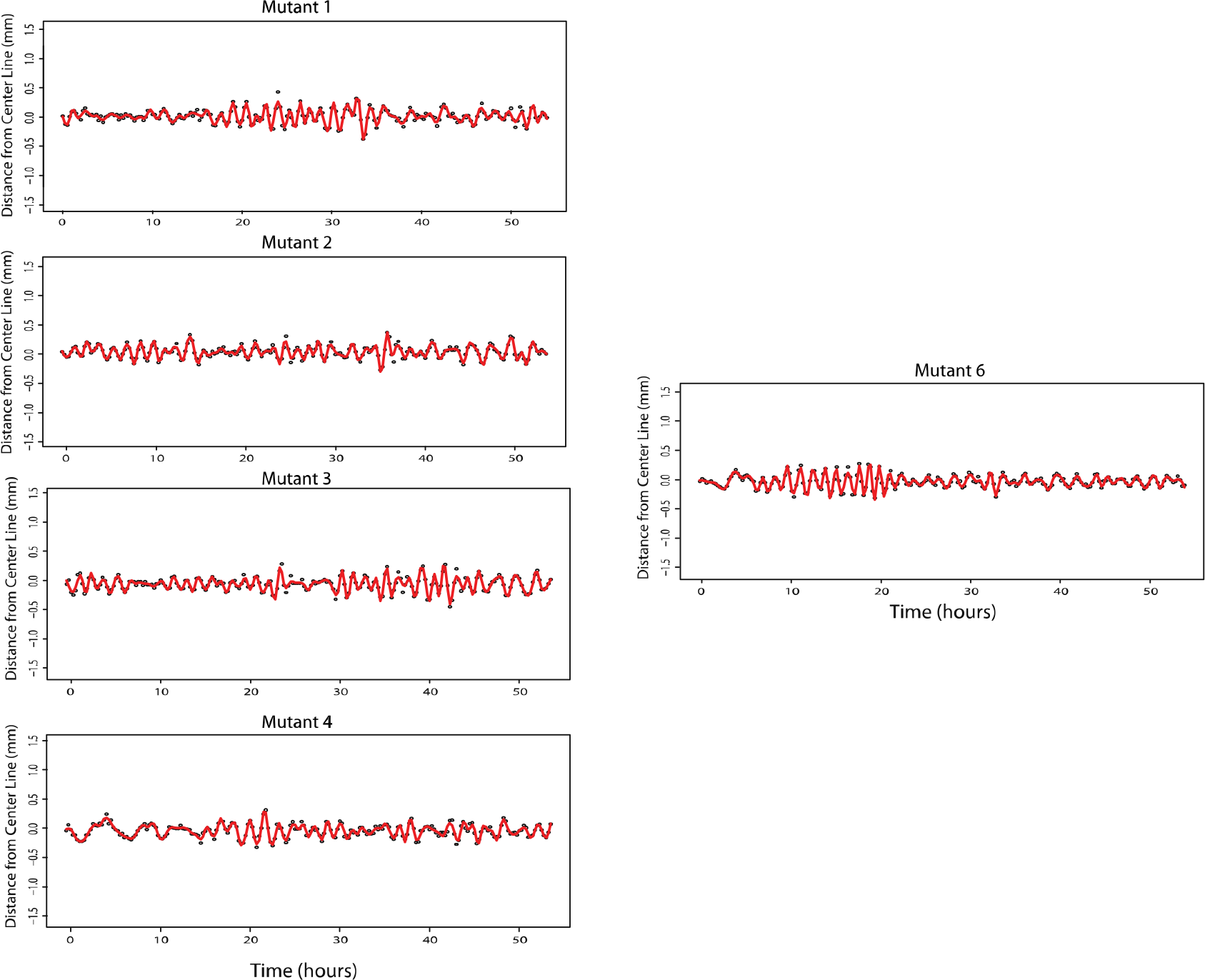
Additional plots of tip position relative to the midline of *oshk1-1* roots throughout 54 hours of growth.

**Extended Figure 8:**
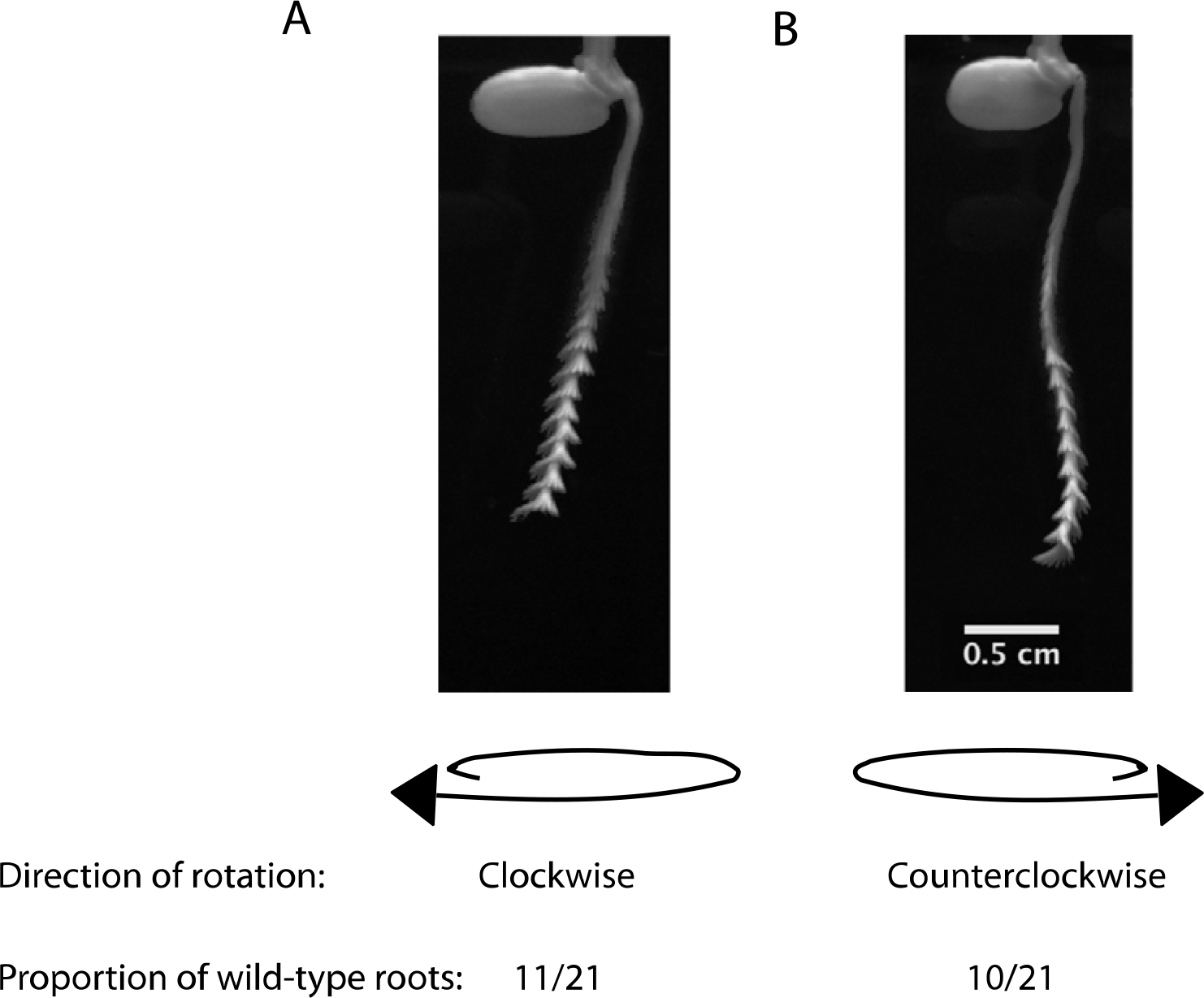
Direction of root rotations inferred from maximum intensity projections of wild-type roots (n=21). Clockwise direction, a, is defined as clockwise movement of the tip as the root moves away from the observer. Counterclockwise direction, b, is defined as counterclockwise movement of the tip as the root moves away from the observer.

**Extended Figure 9:**
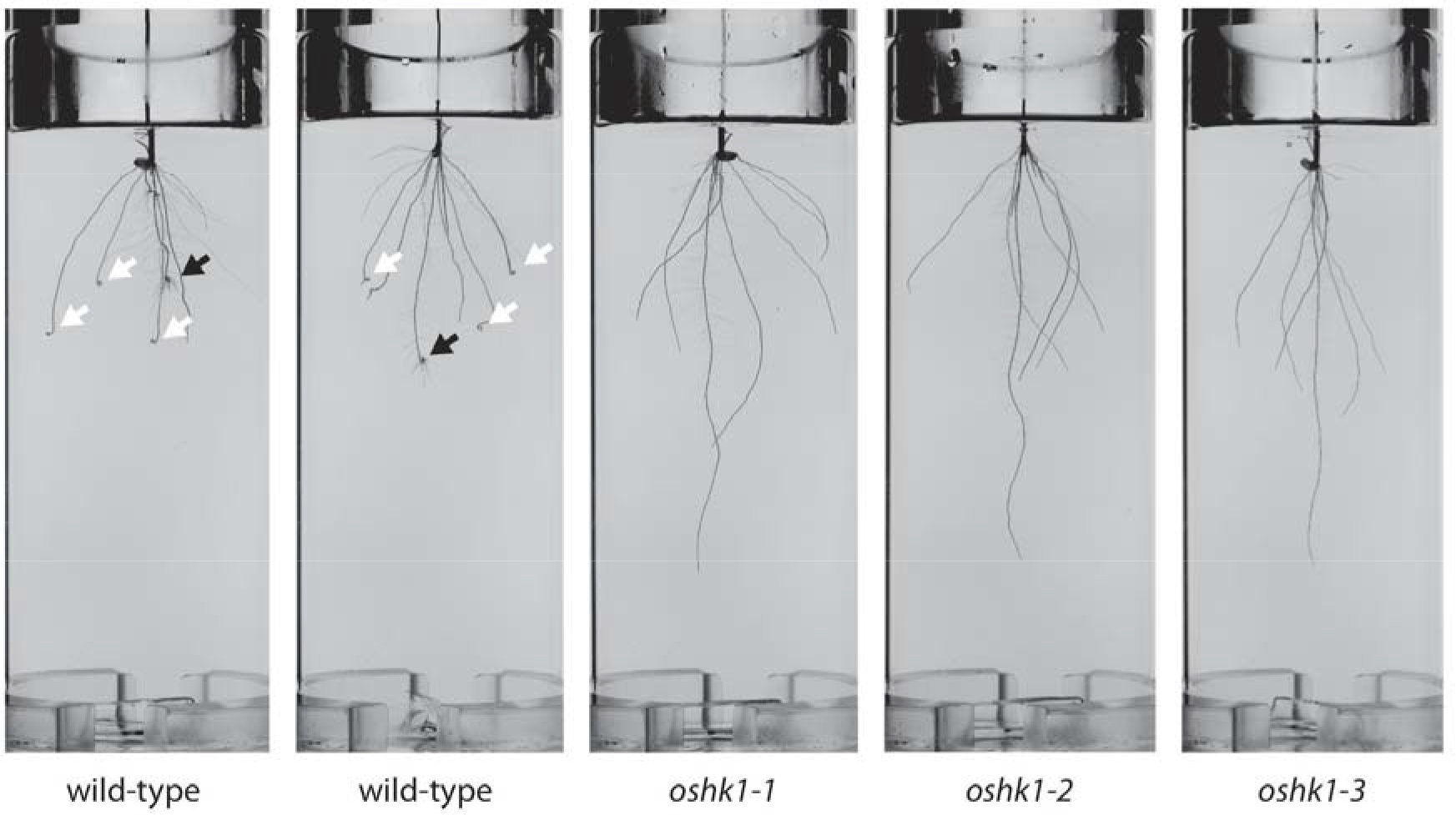
Root meander curling phenotype among wild-type and *oshk1* mutants. Images of seedlings seven days after transplanting into jars with Yoshida’s nutrient solution solidified with 0.25% Gelzan. Black arrows point to curled primary roots, while white arrows show curled crown roots.

**Extended Figure 10:**
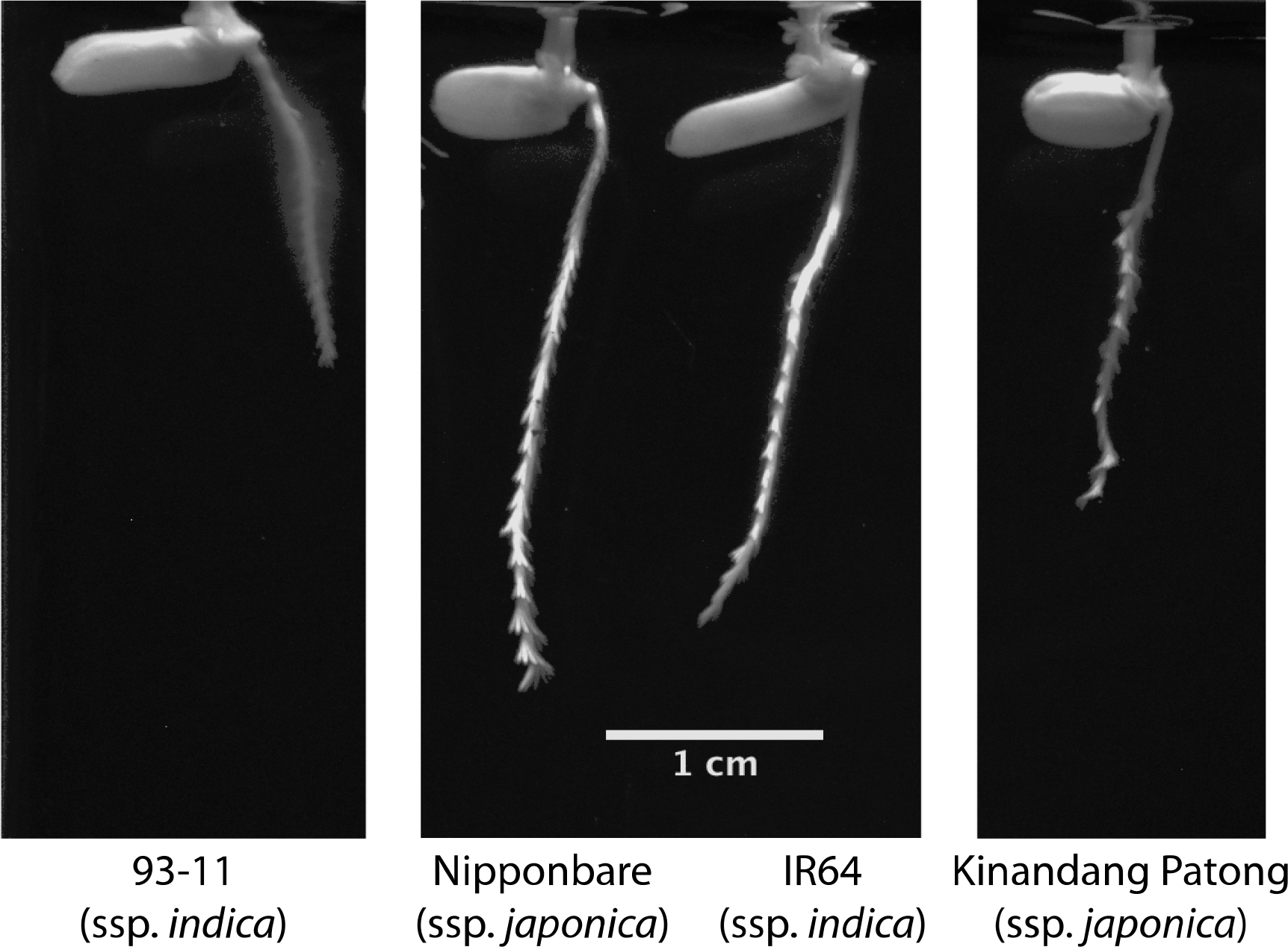
Maximum intensity projections of a diverse set of rice cultivars over 48 hours of growth, imaged at 15-minute intervals.

**Extended Figure 11:**
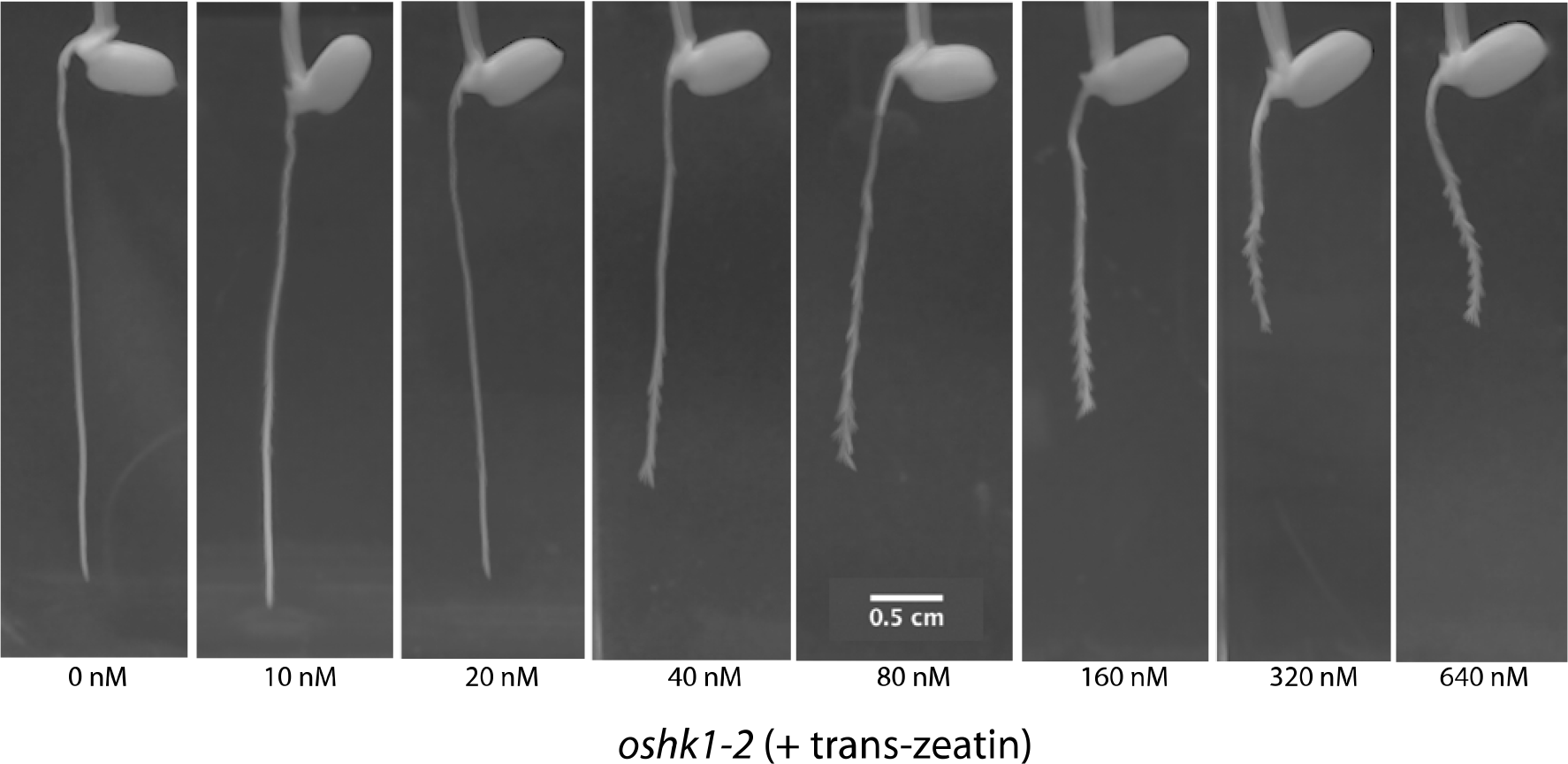
Maximum intensity projections of *oshk1-2* over 54 hours of growth, imaged at 15-minute intervals. Nutrient media, solidified with 0.2% Gelzan, is supplemented with trans-zeatin.

**Extended Figure 12:**
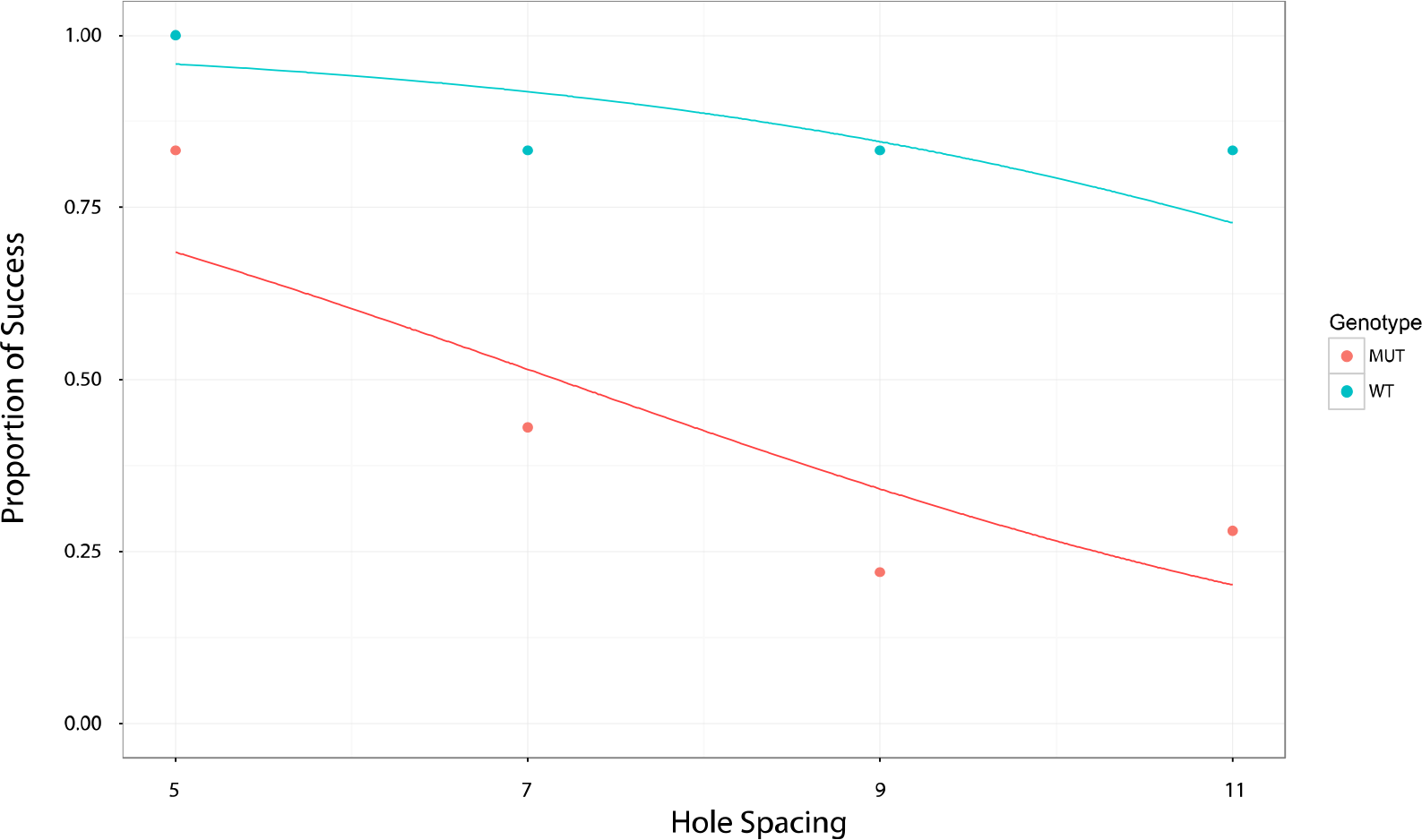
Proportion of success in finding holes in flat platform across 4 different hole spacings. Dots represent observed proportion of successes after 72 hours on surface, curves represent fitted values from logistic regression model.

Supplementary Movie 1: 60 hours of growth of wild-type roots. Images were taken every 15 minutes. Frame rate of the video is 15 fps.

Supplementary Movie 2: 60 hours of growth of *oshk1-2* roots. Images were taken every 15 minutes. Frame rate of the video is 15 fps.

Supplementary Movie 3: 78 hours of growth of wild-type roots, showing in initiation of “Root Meander Curling (RMC)” phenotype. Images were taken every 15 minutes. Frame rate of the video is 15 fps.

## Methods

### Plant Materials and Growth Conditions

All plants are in the X.Kitaake background, a Kitaake line containing the *XA21* gene driven by the maize (Zea mays) ubiquitin promoter ^1^.

Seeds were dehulled, sterilized with bleach, and rinsed. All plant growth was in Yoshida’s nutrient solution solidified with gellan gum (Gelzan, Caisson Inc.).

For RSA experiments, seeds were germinated for two days on petri plates in the dark at 30°C and then transplanted into 2L glass jars containing media solidified with 0.25% Gelzan. Plants were imaged after seven days in a growth chamber at 12hr dark/12hr light cycle and 28°C (day)/26°C (night).

For time-lapse imaging experiments, sterilized seeds were grown directly in GA-7 Magenta vessels containing media solidified with 0.15% Gelzan. After two days in the dark at 30°C, vessels were moved to constant, low light at 24°C for imaging.

For surface exploration assays, sterilized seeds were grown directly in GA-7-3 Magenta vessels containing media solidified with 0.5% Gelzan.

### RSA Imaging and Phenotyping

Plants were imaged and RSA traits measured using the GiaRoots package as in ^2^, except 20 photos per plant were used.

### Identification and Confirmation of OsHK1 Mutants

*oshk1-1* was first identified as a deep-root segregant from the mutant line FN-287. Of the initial six plants phenotyped, three showed this mutant phenotype. All three deep-rooted plants were homozygous for a SBS at position Chr6:26808461, while the other three were either heterozygous or homozygous for the wild-type allele. Primers for genotyping are as follows, FN287_06g44410_F:5’-tccgagcaaaaagaatgatg and FN287_06g44410_R:5’-gcagaaaaactgacagggatg.

*oshk1-2* is from the mutant line FN-179. Of the initial four plants phenotyped, all showed the deep-root mutant phenotype and were homozygous for an inversion between Chr6:25948672-26809852. Primers for genotyping are as follows: Inversion specific, FN179_06g44410_INVF:5’-caggaacagcatgaagaggag and FN179_06g44410_INVR:5’-ttcaggggttcttagcatgg. Wild-type specific, FN179_06g44410_WTF:5’-ttcaggggttcttagcatgg and FN179_06g44410_WTR:5’-catagcccctgaggtaacga.

*oshk1-3* was identified as a deep-root segregant from the mutant line FN790-S. Of the initial six plants phenotyped, three showed this mutant phenotype. All three deep-rooted plants were homozygous for a deletion from Chr6:26807814-26812817, while the other three were either heterozygous or homozygous for the wild-type allele. Primers for genotyping are as follows: Deletion specific, FN790S_06g44410_DELF:5’-aagagctcgttcggaaatca and FN790S_06g44410_DELR:5’-cgctttccttgctttttcat. Wild-type specific, FN790S_06g44410_WTF:5’-aagagctcgttcggaaatca and FN790S_06g44410_WTR:5’-tccataaggcggttcaaatc.

Genomic coordinates are from Os-Nipponbare-Reference-IRGSP-1.0 ^3^

### Quantitative Real-time PCR (qPCR) of OsHK1 *Levels*

Seeds of X.Kitaake were sequentially surface sterilized in 75% ethanol and 50% bleach for 5 minutes before rinsing in sterile water and planting in magenta boxes containing Yoshida’s media. They were placed in a 30 degree incubator in the dark for 2 days before moving to low light conditions at 24 degrees for 2 days. Four 1mm sections from roots at least 1.5 cm in length were harvested starting at the tip and moving shootward. Seven sections were pooled for each sample. A total of 6 replicates for each section were analyzed by qPCR.

RNA was prepared by grinding tissue frozen in liquid nitrogen, using RNAzol RNA isolation reagent according to the manufacturer’s protocol (Sigma-Aldrich). RNA yield was assayed using Qubit RNA quantification system (ThermoFisher Scientific), and RNA integrity was confirmed by agarose gel electrophoresis. First strand cDNA synthesis was performed using the Superscript IV cDNA synthesis kit according to the manufacturer’s protocol (ThermoFisher Scientific). qPCR was performed on a Roche Lightcycler 480 II thermal cycler using FastStart Universal Probe Master Mix (Roche). We identified stably expressed reference genes for rice root qPCR by cross-referencing a list of globally stably expressed genes from ^4^ with genes that exhibited stable expression in a rice root microarray study of crown root developmental zones ^5^. We selected the three reference genes LOC_Os08g18110 (putative alpha-soluble NSF attachment protein), LOC_Os11g26910 (putative SKP1-like protein 1B), and LOC_Os12g41220 (putative ubiquitin-conjugating enzyme) from which we were able to efficiently PCR amplify single bands of the expected size from rice root cDNA.

We analyzed qPCR by taking the geometric mean of the Cp (PCR cycle value) for the three reference genes per sample according to ^6^ and subtracting the Cp value for *OsHK1* to create the delta-Cp measure of transcript abundance. We estimate efficiency of the *OsHK1* qPCR primers by performing 3 biological replicate standard curves across two sequential 5 fold dilutions of rice root cDNA derived from 1mm sections near the root tip and taking the average delta-Cp between serial dilutions. Efficiency was calculated as the ratio of log2(.2) (the dilution factor) to the averaged delta-Cp across the serial dilutions. We then used the estimated efficiency of the primers to transform the delta-Cp measures to obtain an estimate of log2(relative expression level) by taking log2 of the quantity (1+ *OsHKI* primer efficiency)^delta-Cp. We normalized the data to be log2(fold changes) in relation to the lowest average level of expression, which was observed in the 3-4 mm root section, by subtracting the average of log2(expression level) of the 3-4 mm section from each of the sections [see Statistical Methods below].

### Time-lapse Imaging of Root Growth and Quantification of Circumnutation Parameters

Wild-type and mutant seeds of X. Kitaake were surface sterilized and grown in Yoshida’s media containing .15% Gelzan and imaged every 15 minutes for 54 hours using a Marlin machine vision camera (Allied Vision, Inc.). The tip for 9 wild-type and 7 *oshk1-2* plants were manually tracked in ImageJ. We used loess regression to estimate a center line of the root, and calculated the distance in pixels from the center line for each observation to yield a vector of pixel-distance measurements. Pixel distance was converted to an approximate distance in millimeters by scaling to the known width of the visible back of each magenta box using a conversion factor determined by measuring a ruler placed in an empty magenta box in the same plane where seeds were sown. We used a sliding window approach to estimate the amplitude over time by taking the average distance from adjacent maxima and minima from the loess center line. These distances were averaged within bins representing amplitude within 6 hour windows. Period was calculated by measuring the length of time between adjacent maxima or minima (depending on which was observed first in the sequence). These measurements were averaged within bins of 9 hours. An analysis script of this workflow has been uploaded to [Script will be uploaded to Github, available from https://www.dropbox.com/sh/09u9znvxwbvvgnf/AAB-c8ybM3PFyxnfRW2GsVNma?dl=0 during review].

### RNA-Sequencing

3 magenta boxes of wild-type and 1 box each of the 3 allelic mutants were prepared and grown in a manner identical to that for the above qPCR experiment. RNA was isolated from the section 1-2mm from the root tip in a manner identical to the above qPCR experiment. Sections from 12-15 roots from the same box were pooled to form 3 biological replicates for wild-type and 3 for mutant. 150 ng of RNA was input into the Lexogen Quantseq Fwd RNA-Sequencing library kit following manufacturer’s protocol (Lexogen). The 6 libraries were barcoded, pooled, and sequenced on a single lane of Illumina HiSeq 4000. Reads were mapped to the MSU6 build of the rice genome downloaded from https://support.illumina.com/sequencing/sequencing_software/igenome.html. Reads were aligned using hisat2 version 2.0.0-beta with default settings ^7^. Reads mapping to genes were quantified using htseq-count version 0.5.4p3 with default settings utilizing the genes.gtf file downloaded with the MSU6 genome.

### RNA-Sequencing differential expression analysis

RNA-Seq data analysis was performed by importing the counts output by HT-Seq into R for subsequent analysis with the EdgeR package ^8^. We defined “expressed genes” to be those with observed reads in 3 or more libraries. We performed differential expression analysis by comparing the wild-type with mutant using EdgeR commands glmQLFit and glmQLFTest. We defined “differentially expressed” genes to be those with difference between wild-type and mutant with an FDR < .05 and absolute(log2(fold change)) > .4. An analysis script has been uploaded to [Script will be uploaded to Github, available from https://www.dropbox.com/sh/09u9znvxwbvvgnf/AAB-c8ybM3PFyxnfRW2GsVNma?dl=0 during review].

### Cytokinin treatment

Trans-zeatin was prepared in a stock solution of 100 uM in .01 N KOH. We created a dilution series of 0, 10, 20, 40, 80, 160, 320, and 640 nM trans-zeatin. Seeds of the FN-179 mutant were sown in magenta boxes containing Yoshida’s media and .2% Gelzan with varying concentrations of trans-zeatin and imaged using Logitech webcams in a Percival growth chamber under low light conditions at 24 degrees.

### Root exploration assay

Seeds were sown in GA-7-3 Magenta vessels containing media solidified with 0.5% Gelzan. Containers had raised polycarbonate surfaces of equal height with holes of 2.5mm diameter equally spaced in a square grid at distances of 5, 7, 9, and 11mm. Plants were grown under a 12 hour light/dark cycle at 25°C.

An automatic imaging system acquired images of roots from two cameras at 15-minute intervals. A FLIR Flea3 color camera (FL3-U3-13S2C-CS) was positioned facing the front of the growth container to visualize root tip circumnutation and root depth. A second FLIR Flea3 color camera (FL3-U3-120S3C-C) was placed at an angle to capture root growth along the surfaces. The cameras were moved at set intervals by an Arduino controlled horizontal and vertical gantry with stepper motors.

The containers were imaged for a minimum period of five days. A trial was considered a success when the primary root hit a surface and found a hole within 72 hours of first hitting the surface; failures occurred when the root hit the surface but did not find a hole in that time period. Exclusions included trials where the plant had no primary roots, the primary root never hit the surface or directly found the hole without surface contact, the primary root grew out of view of the camera, the primary root exhibited a root meander curling phenotype, or the growth media was visibly contaminated.

### Statistical Methods

All statistical calculations were performed in RStudio Version 1.0.143.

### Root length assay

13, 9, 8, and 8 individual plants were phenotyped for X. Kitaake, FN-179, FN-287, and FN790-S respectively. We utilized pairwise two-sided t-tests assuming equal variance, with the Bonferonni correction for multiple testing. Approximate normality was verified by visually inspecting normal quantile-quantile plots for depth of each genotype. We performed tests of significance at alpha = .05. The p-value for the difference in mean of each mutant line was not significant (p-value = 1 for all 3 comparisons). The p-value for test of each mutant with wildtype was <2e-16. 95% Confidence Intervals (CIs) based on ANOVA for mean of X. Kitaake: (57.13, 66.83), FN-179: (117.97, 125.42), FN-287: (115.30, 126.16), FN790-S: (115.71, 126.58). An analysis script has been uploaded to [Script will be uploaded to Github, available from https://www.dropbox.com/sh/09u9znvxwbvvgnf/AAB-c8ybM3PFyxnfRW2GsVNma?dl=0 during review].

### HK1 qPCR

7 roots of each section were dissected and pooled into 6 individual biological replicates. Individual replicates were grown within single magenta boxes in conditions described above. We defined “expression level” to be (1 + *OsHK1* primer efficiency)^(geometric mean of reference genes’ Cp – *OsHK1* Cp) for each sample. Gene expression is well-known to follow an approximately log-normal distribution^9,10^. Therefore the log of the expression levels is expected to be approximately normally distributed. We normalized the log2(expression levels) to the average of section 4, which exhibited the lowest expression level. Thus, each value can be interpreted as the log2(fold change) of that section compared to the average log2 expression level for section 4. We utilized pairwise two-sided t-tests assuming equal variance, with the Bonferonni correction for multiple testing. Approximate normality was verified by visually inspecting normal quantile-quantile plots for observed log2(expression levels) for each section. We performed tests of significance at alpha = .05. The p-values for the differences in sections 1 vs. section 2 and section 3 vs section 4 were not significant (p-value = 0.2421 and 0.2849, respectively). The p-values for differences in section 1 vs. section 3 and section 1 vs. section 4 were significant (p-value = 0.0095 and 7.3e-05, respectively). The p-values for section 2 vs. section 3 and section 2 vs. section were significant (p-value = 6.1e-05 and 7.6e-07, respectively). 95% CIs based on ANOVA for mean of section 1: (1.13, 1.91), section 2: (1.55, 2.65), section 3: (.01, 1.11), section 4: (-0.55, .55). An analysis script has been uploaded to [Script will be uploaded to Github, available from https://www.dropbox.com/sh/09u9znvxwbvvgnf/AAB-c8ybM3PFyxnfRW2GsVNma?dl=0 during review].

### Cytokinin response gene enrichment analysis

We performed over-enrichment analysis by taking the intersection of the lists of differentially expressed genes identified in this study (higher and lower in wildtype compared to mutant, respectively) with the lists of genes induced and repressed by cytokinin identified in the whole root study of ^11^. We compared the number of genes in the intersection of the wild-type high/cytokinin induced sets and the wild-type low/cytokinin repressed sets with the expected number of genes in the intersection of each respective comparison assuming independence of set membership. We utilized a one-sided hypergeometric test to calculate p-values (p-value = 0 and 1.93e-08 for test of over-enrichment, respectively). An analysis script has been uploaded to [Script will be uploaded to Github, available from https://www.dropbox.com/sh/09u9znvxwbvvgnf/AAB-c8ybM3PFyxnfRW2GsVNma?dl=0 during review].

### Gene Ontology analysis

We uploaded lists of differentially expressed genes to the Agrigo V2 server ^12^. We analyzed the gene sets compared to MSU7 gene annotations, using the Fisher method, and Yekuteli FDR < .05.

### Logistic regression of hole finding probability

11, 12, 6, and 6 roots of wildtype and 12, 16, 18, and 7 roots of mutant were allowed to grow on a platform for 72 hours at hole spacings of 5mm, 7mm, 9mm, and 11mm respectively. “Success” was defined as the root tip encountering and growing into a hole. We modelled the probability of success using logistic regression with hole spacing and genotype as covariates. Based on the Wald test, we observed significant effects of both genotype and spacing at alpha of .05 (p-value = 0.000211 and 0.007853, respectively).

We utilized profile likelihood based Confidence Interval estimation to determine confidence intervals for the odds ratios (95% CI for Odds Ratio for Genotype: (3.35, 42.63), 95% CI for spacing: (0.53, 0.90). An analysis script has been uploaded to [Script will be uploaded to Github, available from https://www.dropbox.com/sh/09u9znvxwbvvgnf/AAB-c8ybM3PFyxnfRW2GsVNma?dl=0 during review].

